## Supplementary material for "Rheb membrane orientation dynamics and functional consequences elucidated by molecular simulations, single-molecule-FRET and signaling assays": SI Figures and Text

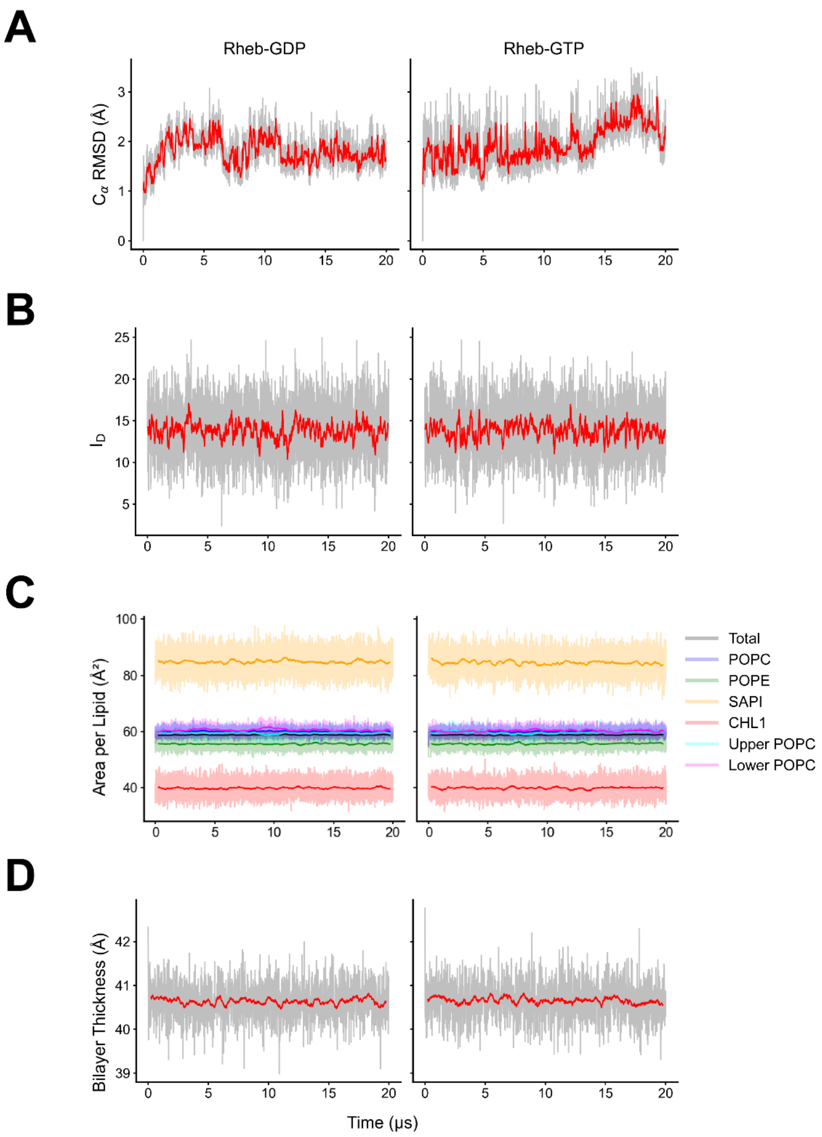


**Figure S1: Time-evolution of protein and membrane structures during Rheb-GDP (left) and Rheb-GTP (right) simulations.** (**A**) $C_\alpha$ atoms RMSD of G-domain residues 1-170 from the equilibrated snapshot excluding the flexible switch I (residues 33-41) and switch II (residues 63-79). (**B**) Insertion depth ($I_D$) of the farnesyl tail into the bilayer core, defined as the z-distance between the centers-of-mass (COM) of the membrane and the farnesyl acyl chain. (**C**) Area per lipid of each lipid type computed as described recently[1]. (**D**) Membrane thickness measured as the z-distance between upper and lower leaflet phosphate atoms COM. Data sampled every 1 ns are shown as semitransparent lines with 50 ns moving averages in heavier solid lines.

**SI Movie 1**: 10 μs excerpts from the production Rheb-GDP and -GTP membrane simulations. Lipid phosphate atoms are shown as slate spheres, and Rheb is rendered as a light-gray cartoon, anchored to the membrane by its farnesylated C-terminus (cyan sticks). Switch I (residues 33–41, pink), switch II (residues 63–76, teal), α3 (residues 90–107), α4 (residues 131–140, yellow), and α5 (residues 153–170, orange) are highlighted to indicate the orientation of the G-domain relative to the membrane during the trajectory. Bound GDP is shown as green transparent sticks. snapshots were rendered every 5 simulation frames (2 ns) and temporally interpolated to generate smoother motion for visualization only.

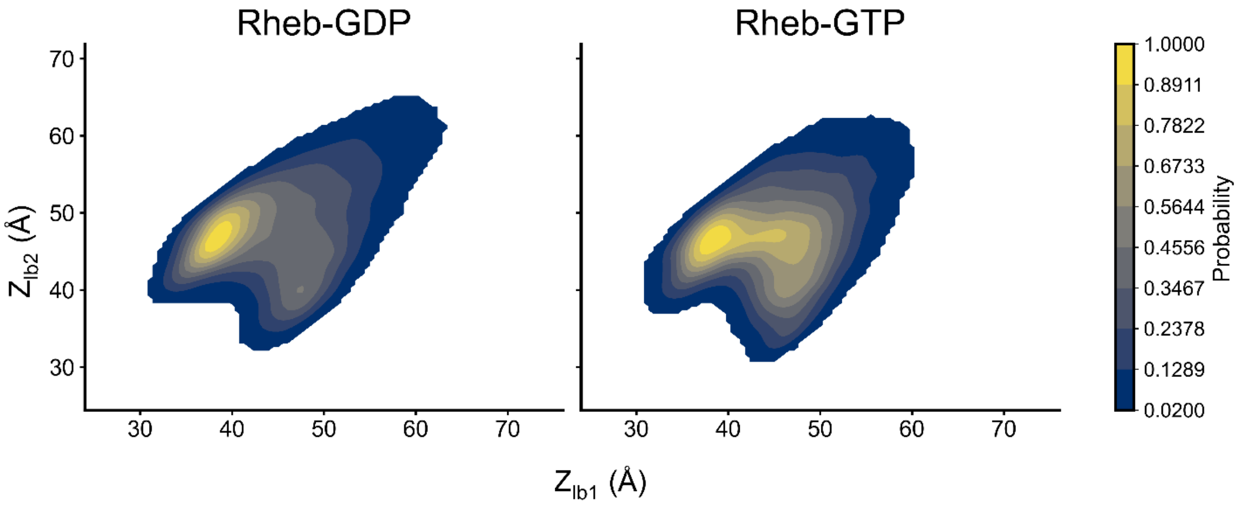


**Figure S2: G-domain dynamics based on previously used reaction coordinates.** Joint distribution of the z-distances of lobe 1 ($Z_{lb1}$; residues 1–85) and lobe 2 ($Z_{lb2}$; residues 86–170) from the bilayer center. Distances were computed as the difference in z-coordinates between the COMs of the bilayer and backbone atoms of each lobe and normalized by the maximum value for visualization. The Rheb–GTP distribution recapitulates a previous report[2].

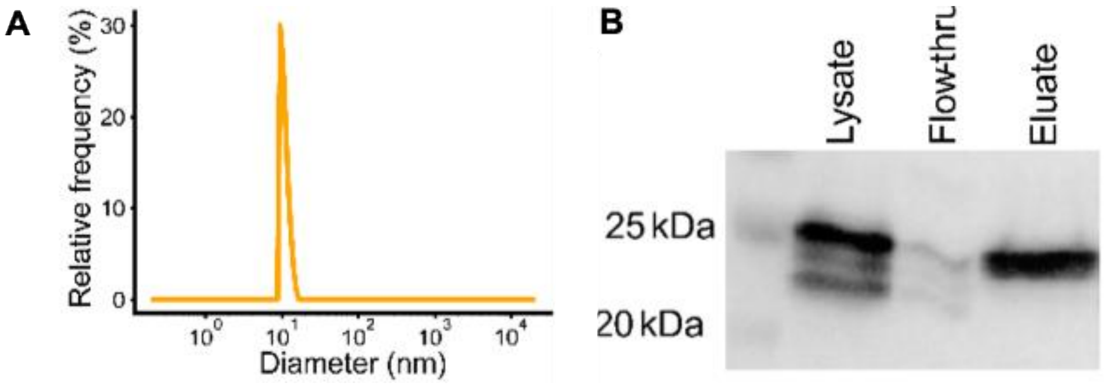


**Figure S3: Isolation of Rheb in cell-derived lipid nanodiscs.** (**A**) Histogram of particle diameters from 16 independent DLS measurements, showing sizes within the expected range and indicating a successful purification and homogeneous nanodisc population. (**B**) Western blot showing levels of Rheb in SMA-treated lysate, flow through after HA-tag binding, and the HA-bead eluate. We can see that our HA-tagged Rheb is expressed and present in the eluate.

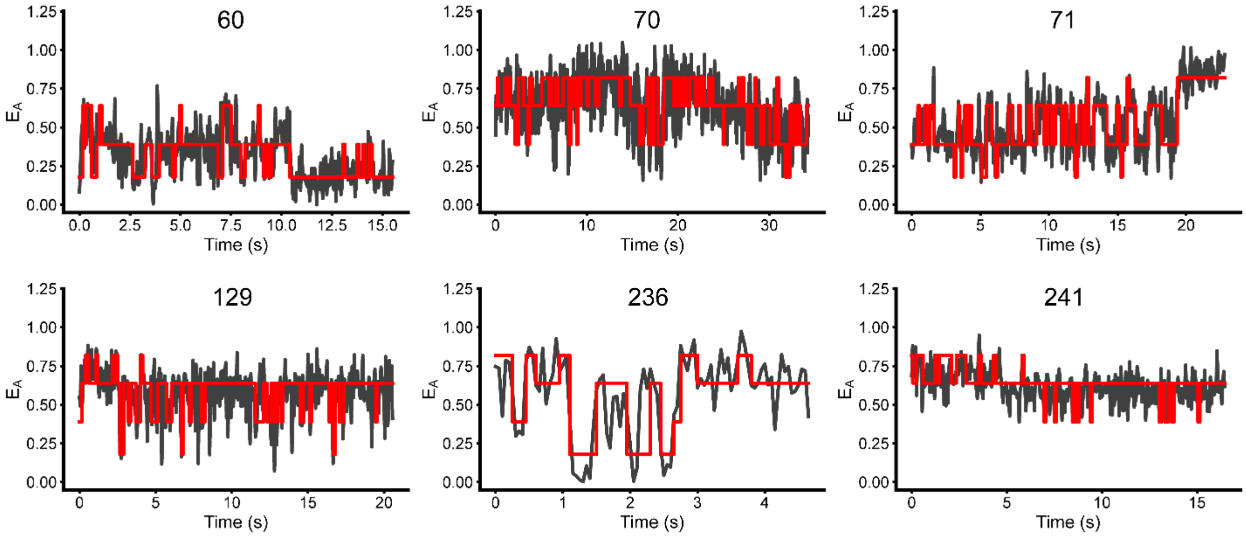


**Figure S4: Representative smFRET traces of Rheb in nanodiscs.** The raw fluorescence efficiency ($E_A$) is shown together with a discrete state trajectory (red) obtained by assigning each frame to the nearest experimentally determined smFRET state mean (main Fig 3B) after smoothing with a 3-frame running window. These traces provide an approximate visualization of state transitions but do not represent an independent hidden Markov or STaSI idealized trajectory.

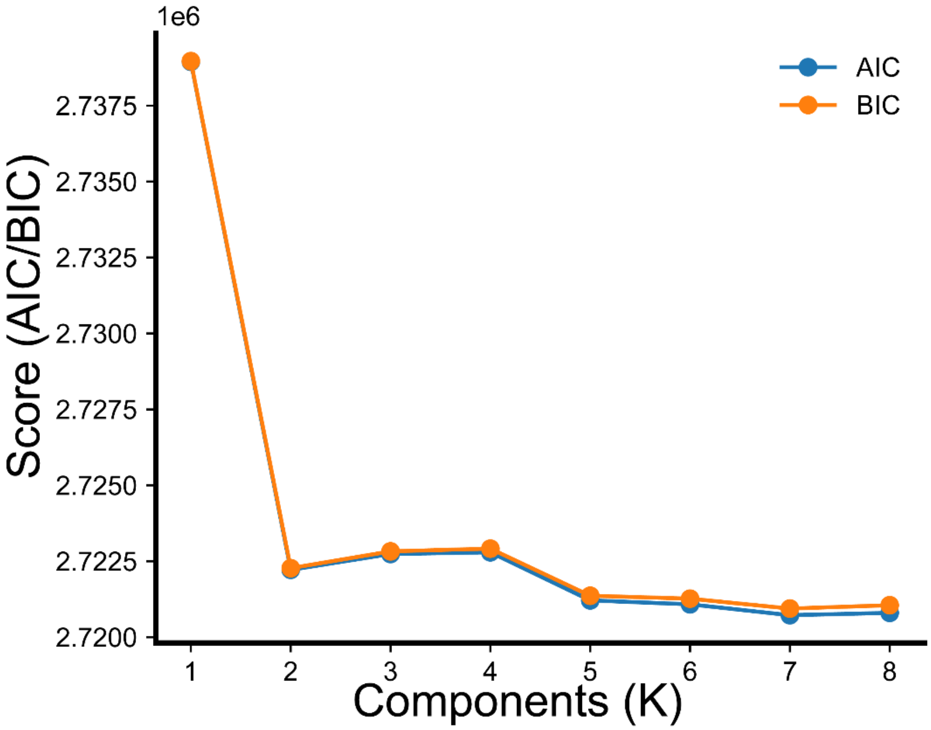


**Figure S5:** Akaike information criterion (AIC) and Bayesian information criterion (BIC) values for Gaussian mixture models fitted to the unbinned $R_{eff}$ distance distribution as a function of the number of mixture components (k). Both criteria decrease sharply from k = 1 to k = 2 and show only marginal improvement beyond k = 3–4, indicating an early saturation of model complexity. A four-component model was therefore selected as a representation consistent with the number of smFRET-derived states.

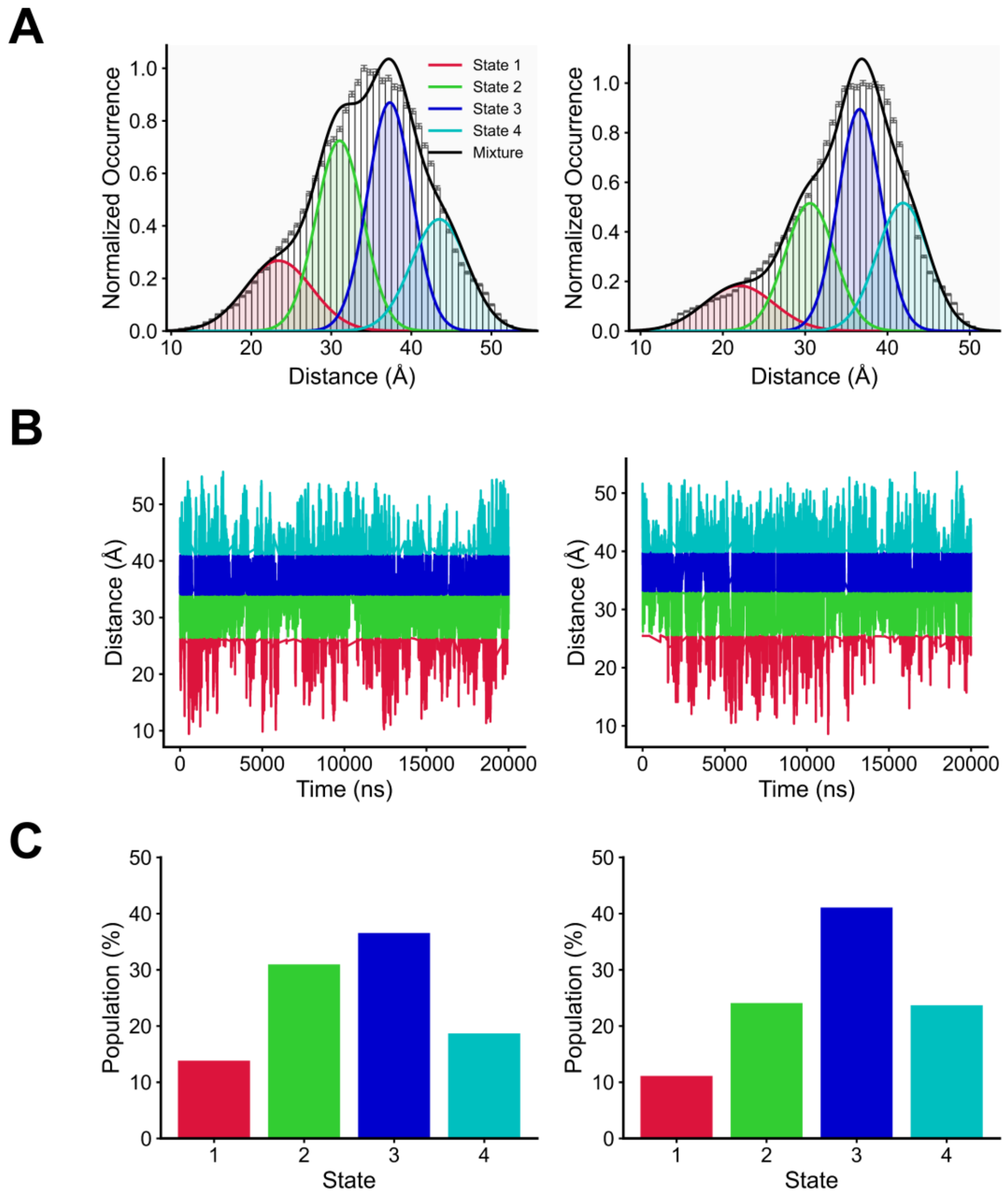


**Figure S6: Conformations sampled in Rheb-GDP (left) and Rheb-GTP (right) simulations.** (**A**) Histograms of $R_{C\alpha}$ with a four-component Gaussian mixture model (GMM) decomposition overlaid (colored curves) with the total mixture fit shown in black. (**B**) $R_{C\alpha}$ time series with frames colored by assigned GMM states obtained by hard classification to the nearest GMM component mean (equivalently, maximum-likelihood component assignment for well-separated components). (**C**) GMM state populations computed from the corresponding state assignments.

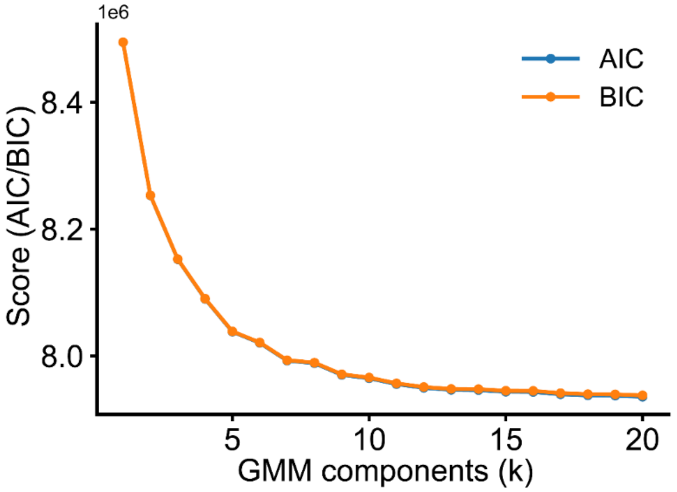


**Figure S7: AIC/BIC-based selection of GMM complexity for spherical density estimation.** Akaike information criterion (AIC) and Bayesian information criterion (BIC) scores were computed for the GMM fit with *k* components to the spherical orientation distribution. Lower values indicate a better tradeoff between goodness-of-fit and model complexity. Both criteria decrease steeply from small *k* and then show diminishing improvement beyond ~10 components, supporting the use of a 10-component GMM for subsequent spherical probability density function (PDF) estimation.

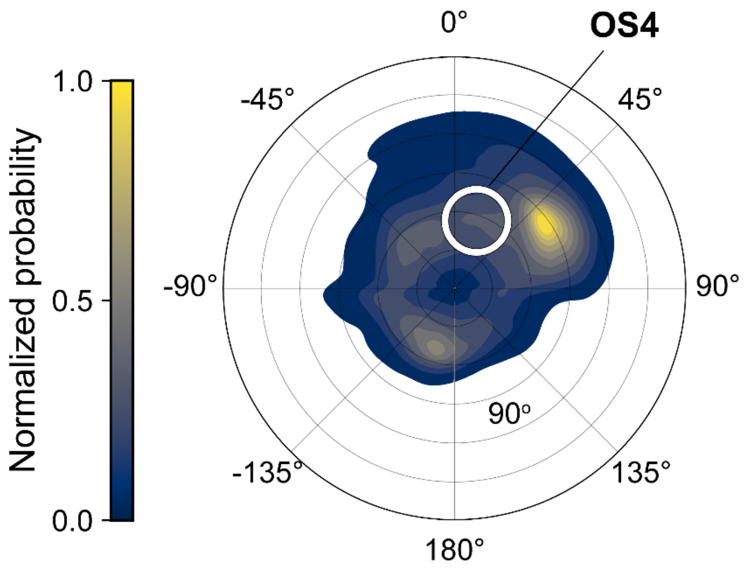


**Figure S9**: **2D probability density obscures some states.** Normalized joint probability distribution of θ and φ computed from the combined GDP- and GTP-Rheb trajectories. The annotated circle marks the angular location of OS4, which lies within a contiguous shoulder of the dominant density region. This overlap illustrates how a (θ, φ) analysis may obscure low-population states, motivating the use of an additional orthogonal coordinate ($R_{eff}$) for further classification of conformers.

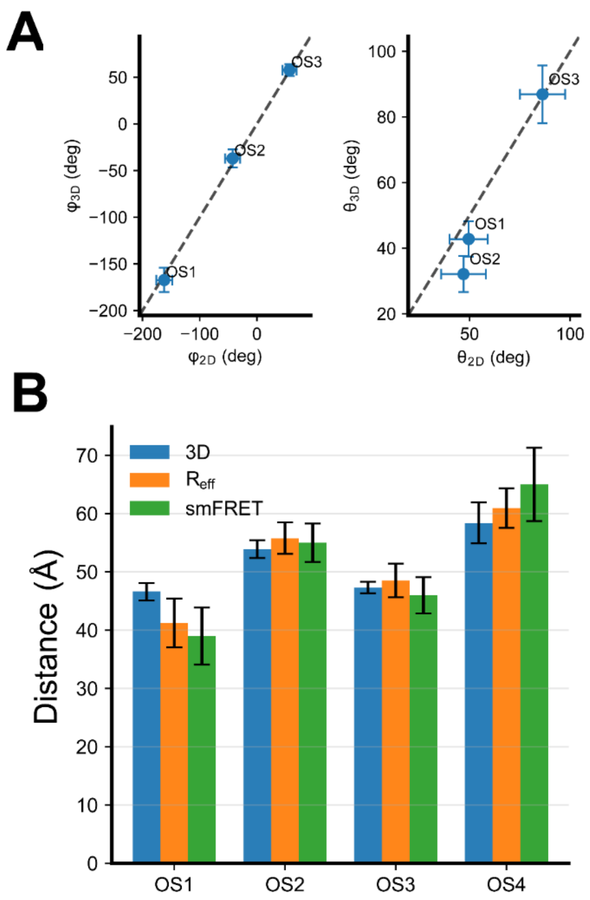


**Figure S9: Classification of simulated conformers using different RCs yields similar pools of ensembles.** (**A**) Scatter plots comparing tilt and rotation angles in orientations states derived from 2D (θ, φ) and 3D ($R_{eff}$, θ, φ) analyses of the combined GDP- and GTP-Rheb trajectories. Points represent the geometric center of ensemble distributions with horizontal and vertical error bars representing SDs in 2D- and 3D-derived ensembles, respectively. OS4 was omitted because it was not resolvable in the 2D analysis. (**B**) 3D embedding (blue) and $R_{eff}$ (orange) recapitulate distance profiles observed in smFRET (green). smFRET efficiencies were converted to distances and experimental states were assigned to the corresponding orientation states based on the closest mean inter-residue distance. OS1 and OS3 correspond to the intermediate FRET distance states 1 (42.2 Å) and 2 (49.2 Å) or to mean $R_{eff}$ values of 44.0 and 45.0 Å, respectively; OS2 corresponds to the moderately extended smFRET state 3 ($R_{eff}$ 54.0 Å) and OS4 to FRET state 4.

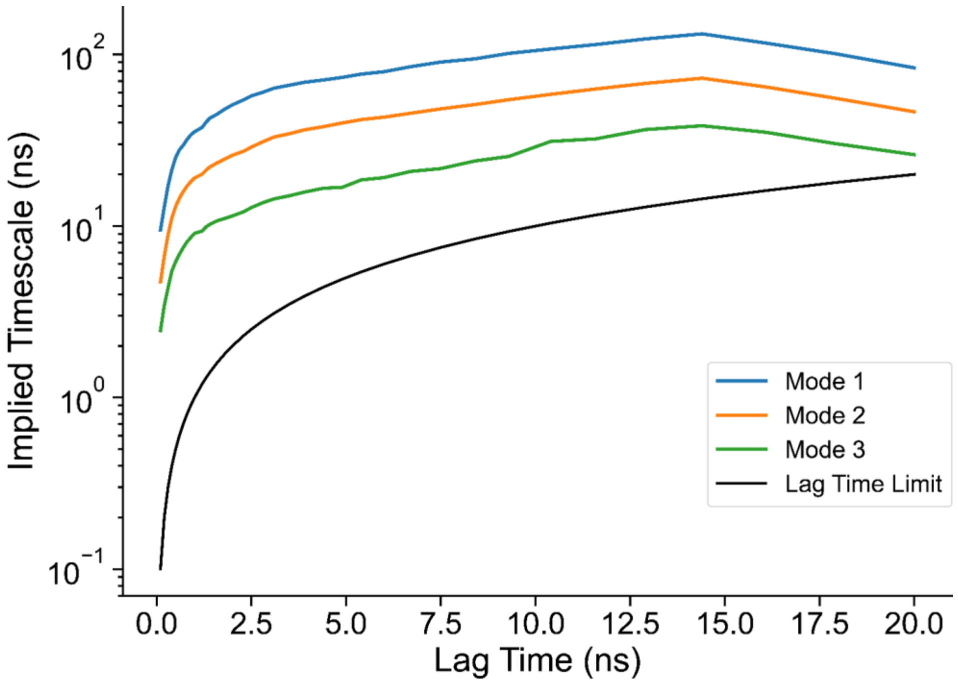


**Figure S10: Implied timescales for Bayesian HMM.** Implied relaxation timescales are shown for the three slowest dynamical modes as a function of lag time. The black curve indicates the lag-time limit (τ = lag). Convergence of the implied timescales at longer lag times indicates approximate Markovian behavior.

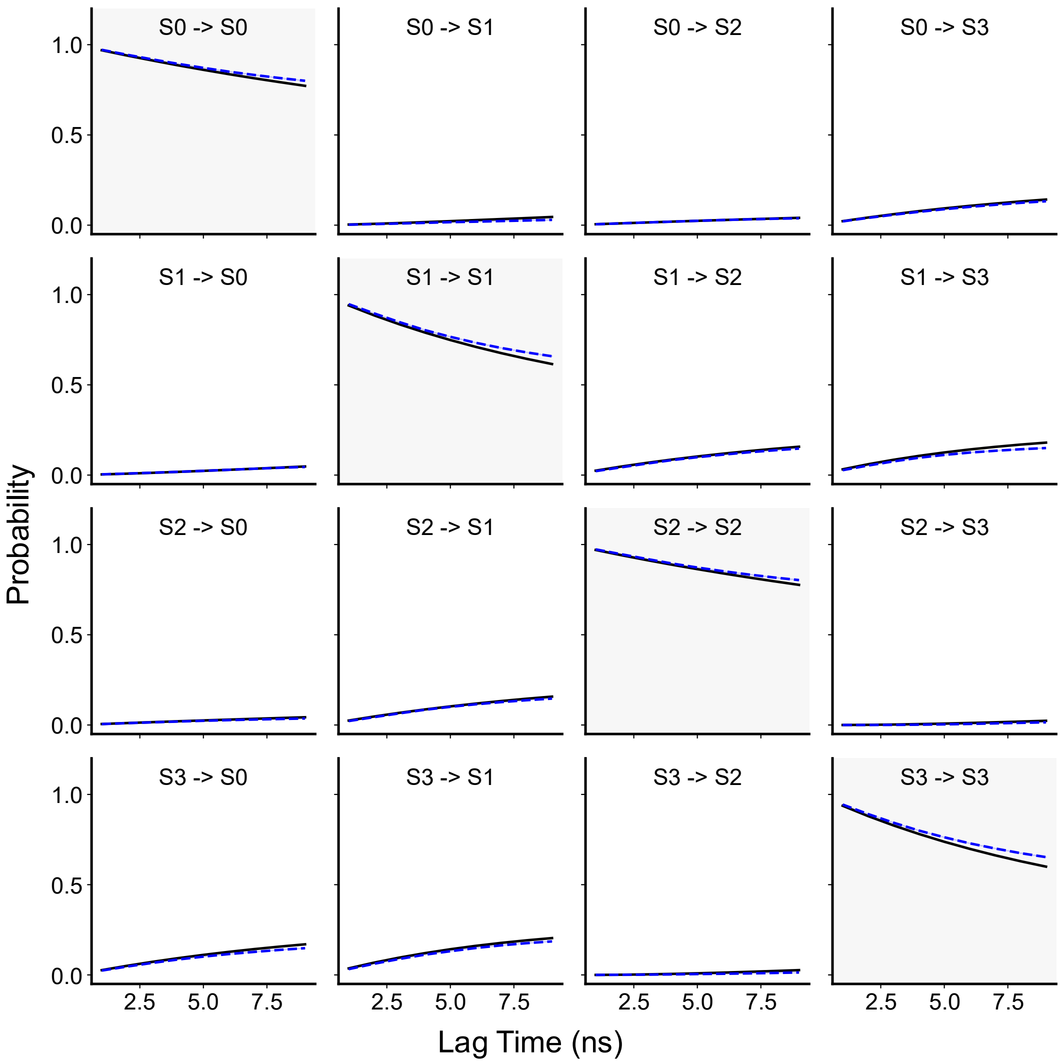


**Figure S11:** Chapman–Kolmogorov (CK) test for HMM validation. For each state pair ($S_i \rightarrow S_j$), transition probabilities are shown as a function of lag time. Solid lines indicate HMM-predicted transition probabilities (propagated from the estimated transition matrix), and dashed lines indicate transition probabilities estimated directly from the simulated trajectories. Agreement between predicted and observed probabilities across lag times supports the Markovian predictive power of the HMM at the selected lag time.
